# Trapped Ion Mobility Spectrometry Reduces Spectral Complexity in Mass Spectrometry Based Workflow

**DOI:** 10.1101/2021.04.01.438072

**Authors:** Joshua Charkow, Hannes L. Röst

## Abstract

In bottom-up mass spectrometry based proteomics, deep proteome coverage is limited by high cofragmentation rates. This occurs when more than one analyte is isolated by the quadrupole and the subsequent fragmentation event produces fragment ions of heterogeneous origin. One strategy to reduce cofragmentation rates is through effective peptide separation techniques such as chromatographic separation and, the more recently popularized, ion mobility (IM) spectrometry which separates peptides by their collisional cross section. Here we investigate the capability of the Trapped Ion Mobility Spectrometry (TIMS) device to effectively separate peptide ions and quantify the separation power of the TIMS device in the context of a Parallel Accumulation-Serial Fragmentation (PASEF) workflow. We found that TIMS IM separation increases the number of interference-free MS1 features 9.2-fold, while decreasing the average peptide density in precursor spectra 6.5 fold. In a Data Dependent Acquisition (DDA) PASEF workflow, IM separation increased the number of spectra without cofragmentation by a factor of 4.1 and the number of high quality spectra 17-fold. This observed decrease in spectral complexity results in a substantial increase in peptide identification rates when using our data-driven model. In the context of a Data Independent Acquisition (DIA), the reduction in spectral complexity resulting from IM separation is estimated to be equivalent to a 4-fold decrease in isolation window width (from 25Da to 6.5Da). Our study shows that TIMS IM separation dramatically reduces cofragmentation rates leading to an increase in peptide identification rates.

## Introduction

Mass spectrometry (MS) based proteomics enables high throughput quantification and identification of thousands of proteins and can provide insights into global function of biological systems. One of the most frequently used approaches is the bottom up proteomics workflow with data dependent acquisition (DDA). In this approach, trypsinized peptides are first separated by liquid chromatography (LC) and as the peptides flow out of the column, they are ionized and transported to the mass analyzer. First, a MS1 spectrum is captured measuring the

mass-to-charge (m/z) and intensity of all incoming ions. Next the ion beam is filtered in the m/z dimension to the chosen precursor based on the previously acquired MS1 spectrum. The filtered ion beam is fragmented and the corresponding MS2 spectrum is then captured. Peptides can be identified using the m/z derived from the MS1 spectra and the fragment ion series captured in the MS2 spectra.

With increasing speed and sensitivity, instruments are able to sequence precursor ions of ever lower intensities, which makes it increasingly hard to obtain high quality fragment ion spectra without cofragmentation. Cofragmentation occurs when two or more peptides are coisolated by the quadrupole selection in the same MS2 spectrum resulting in a chimeric spectrum. Chimeric spectra have a detrimental effect on peptide identification rates ^1^ and cause ratio compression in isobaric labelling workflows, reducing peptide quantification, precision, and accuracy.^2–6^ While the identification issue can be partially addressed computationally,^7,8^ chimeric spectra also challenge the limited intra-scan dynamic range of current mass analyzers, which leads to low-quality ions being recorded for the lower abundant peptide and often only the dominant peptide can be identified with high certainty.^9^ It is therefore preferable to isolate pure precursors, by performing off-line peptide separation ^2^ or through an additional step of isolation using a quadrupole mass filter as in MS3-based workflows.^10^

However, these workflows drastically increase analysis time and cost. Alternatively, on-line separation using ion mobility (IM), which separates analytes by their collisional cross section, has been proposed as a partially orthogonal peptide separation technique to mass spectrometry. IM-MS workflows have been shown to increase dynamic range,^11,12^ proteome coverage ^13,14^ and quantification accuracy.^15–18^ Wide-spread adoption of on-line IM separation was previously hampered by reduced ion transmission and cumbersome geometry, ^17,19,20^ often requiring very long ion paths to achieve the desired resolution.

However, the recent introduction of Trapped Ion Mobility Spectrometry (TIMS) overcomes some of these limitations, providing high ion transmission within a small geometry (5-10 cm). This instrument traps ions at different positions using the opposing forces of gas flow and an electric field ion, separating ions by their collisional cross section, followed by sequential release.^21,22^ The TIMS device supports simultaneous ion accumulation and IM separation resulting in a high duty cycle.^23^ Recently, TIMS has been utilized in a novel acquisition workflow named Parallel Accumulation-Serial Fragmentation (PASEF).^19,24^ Briefly, PASEF involves synchronizing the quadrupole isolation window with the elution of ions from the TIMS device so that multiple precursors can be isolated and easily resolved in a single ion mobility elution cycle. PASEF has been reported to increase sequencing speed 4-10 fold without a loss of sensitivity (sensitivity =∼95%) resulting in high peptide identification rates.^19,24^ PASEF was first applied in a DDA workflow where over 6000 proteins were identified from a HeLa cell digest in a single run.^24^ More recently, PASEF was implemented in a SWATH Data Independent Acquisition (DIA) workflow ^25^ resulting in the identification of over 7600 proteins in a single run.^26^

Although it is clear that the PASEF acquisition workflow achieves high peptide and protein identification rates, the underlying mechanisms contributing to the observed increase in identifications is poorly understood. Specifically, the PASEF sampling scheme results in improvements to ion focusing, increased sequencing speed, as well as the addition of ion trapping and IM separation. Since all of these factors are intertwined, it has so far been difficult to distinguish their impacts on the improvements observed in a PASEF workflow. In this study, we aim to quantify the influence of one of these factors, TIMS IM separation, independently from the other components of the PASEF workflow. Separation power is evaluated by comparing peptide density, cofragmentation rates, precursor ion fraction (PIF), and identification rates of PASEF-experimental data ^24^ simulated with and without IM separation. Furthermore, the degree of cofragmentation and identification rates was also evaluated in the context of a DIA PASEF experiment. We show that application of TIMS IM separation confers substantial benefits to both DDA and DIA workflows which are due to a drastic reduction of peptide density and cofragmentation rates resulting in increased peptide identification rates.

## Results

### TIMS IM separation decreases feature density

To investigate the separation power of TIMS IM, we examined the impact IM separation would have on the density of features which are defined as isotopic envelopes across the IM and retention time (RT) dimensions. Features were identified by MaxQuant from a tryptic HeLa cell digest, acquired on a timsTOF Pro (Bruker) instrument in DDA-PASEF mode.^24,27^ For the 401 923 features identified by MaxQuant, we computed the pairwise overlap matrix (all-against-all) and for each feature, recorded the number of overlapping features. We found that IM separation increased the proportion of features without any overlap 9.2-fold, from 5% to 46% (Figure 1A). The impact of IM separation is illustrated by comparing a representative region with and without IM separation. Without IM separation (Figure 1C), the fundamental difficulty of isolating a pure peptide analyte from a very dense peptide space becomes evident: within a 15 second region in RT there are eleven highly overlapping isotopic envelopes in a 8 m/z region. We can clearly see how the addition of the IM dimension greatly increases separation power by reducing feature overlap and completely resolving all overlaps of half of the identified peptides shown (Figure 1D).

**Figure 1:**
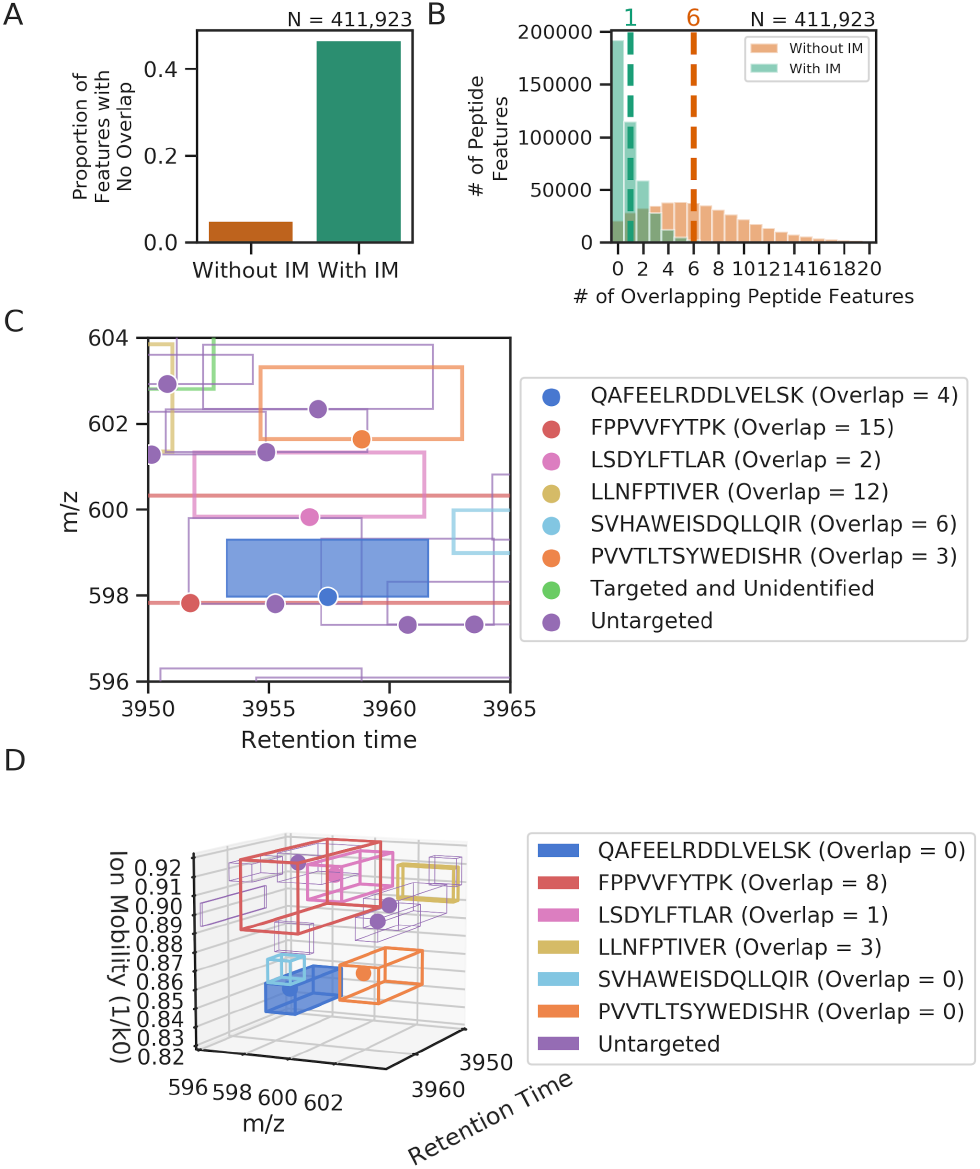
Ion mobility reduces feature density. (A) Proportion of peptide features with no interference with (green) and without (orange) and IM. IM increases the proportion of features without interference 9.2-fold. (B) Histogram of features with (green) and without (orange) IM binned by their degree of interference. Dashed lines indicate the median of each distribution. On average, IM features have 6.5-fold less pairwise overlaps compared to features without IM. (C) Model of peptide features in a representative region of the retention time and m/z space. Peptide features are colored by their identification. Each rectangle spans the isotopic envelope and elution of the peptide and each circle represents the peak apex and monoisotopic m/z of the feature. The entire elution profile for all of the features is not shown and thus not all of the overlaps are shown for every feature. (D) Model of the same region as figure (C) with the IM dimension and peptide IM elution profile added. Roughly 10% of the IM range is shown and thus not all peptide features from figure B are visible. IM reduces the number of overlapping peptide features for all features shown and resolves all interferences in the QAFEELRDDLVELSK (dark blue) peptide.

By computing the number of overlaps per feature with and without IM, we estimated the impact of IM on potential cofragmentation events. We found that the IM dimension reduced the median number of overlaps 6-fold (from 6 to 1), drastically shifting the overlap distribution towards 0 (Figure 1B), and reduced the total number of pairwise overlaps 6.5 fold (Supplemental Figure 1). This observed reduction in feature overlap highlights the degree that IM separation simplifies the feature space by resolving previously interfering signals, and decreasing potential cofragmentation events.

Although the average feature benefits 6-fold from IM separation, we reasoned that the influence of IM separation may depend on feature density, providing a larger benefit in dense regions. By comparing IM separation power in feature dense and sparse regions we found that IM separation is most impactful at high feature density regions. In high feature density regions, the reduction in feature density due to IM is more then 2-fold higher than in the least dense regions (Supplemental figure 2). This substantial variance in feature density reduction indicates that IM separation provides the strongest benefit in samples of high complexity.

To contrast the IM separation power to m/z and RT separation power, we projected each peptide feature to a single dimension and computed pairwise overlaps across a single dimension. We found that, on average, features projected to the m/z and RT axes contained less overlaps compared to features projected in the IM dimension (Supplemental Figure 3).

These results suggest that the peak capacity for IM is lower than RT and m/z peak capacity. Although IM peak capacity may be lower compared to RT or m/z, coupling IM to the mass spectrometry workflow provides a considerable reduction to possible cofragmentation events and observed peptide interferences.

### TIMS IM decreases spectral complexity in a DDA-PASEF workflow

Coupling TIMS IM to MS enables isolation of precursors in both the m/z and IM dimensions simultaneously rather than solely in the m/z space. To investigate whether IM enhances isolation window selectivity, we measured the effects of IM on the rates of cofragmentation. Peptide cofragmentation occurs when two or more peptides are isolated and fragmented during a quadrupole isolation cycle, thus leading to a chimeric MS2 spectrum which has a lower likelihood of correct identification.^1^ We computed cofragmentation rates based on a acquired PASEF run from HeLa cell digest by determining the proportion of windows isolating a single peptide feature with and without IM. IM enhanced isolation windows were directly extracted from the experimental data and non-IM enhanced isolation windows were constructed by collapsing the ion mobility axis, thus considering all features within the m/z isolation window independent of their IM. It is important to note that this does not take into account the lower sequencing speed of an instrument without IM separation, thus making our estimation highly conservative. We found that the number of quadrupole isolation windows containing only a single, pure precursor was 4.1-fold greater (from 14.2% to 59.0%) than the equivalent windows without IM separation. Furthermore, the number of MS2 spectra with at least one cofragmented peptide decreased by 2.1-fold (from 85.8% to 41.0%) with IM separation (Figure 2A). These results also indicate that the majority of MS2 spectra with IM only isolate a single feature (59.0%) in contrast to MS2 spectra without IM where most spectra are chimeric and isolate multiple features (85.8%). To further quantify the effect of TIMS IM on MS2 spectral complexity, the number of features that were isolated in each MS2 spectrum was computed. In our data, an IM enhanced MS2 spectrum contained an average of 2.6-fold fewer features per spectrum compared to equivalent spectra without IM separation, reducing the mean number of isolated features from 4.2 to only 1.6. (Figure 2B). The observed reduction in the rate and degree of cofragmentation resulting from IM isolation, suggest that TIMS IM reduces spectral complexity in a DDA PASEF workflow and yields more pure precursors for MS2 analysis.

While computing the number of coisolated features provides insight into the scope of the problem, our approach so far does not take into account peptide intensity. It is important to consider peptide intensity because this property affects the nature of the cofragmentation event. For example, if a high intensity precursor was cofragmented with a low intensity precursor, the resulting MS2 spectrum may mainly contain signals from the high intensity precursor and will behave differently in downstream analysis compared to a spectrum with cofragmentation of two equally intense precursors. To address this peptide intensity, we computed the precursor ion fraction (PIF) for each isolation window with and without IM in the simulated instrument acquisition. PIF measures the proportion of the isolated ion beam that can be attributed to the targeted precursor. Based on our analysis and a previous study examining the rate of peptide identification as a function of PIF,^9^ we classified isolation windows as either low, standard or high quality based on their PIF value. Using IM separation increases the number of standard quality isolation events 4.1-fold (from 11.2% to 46.1%) and decreases the number of low quality isolation events 3.0-fold (from 87.3% to 29.3%) (Figure 2C). Interestingly, using IM separation improves the average spectrum from containing less than 17% of the ion current from the targeted precursors (with the other 84% contributed by an average of 3.2 other ion species) to containing more than 50% of ion current from the targeted precursor (Supplemental Figure 4). The increase in PIF associated with IM separation suggests that the addition of TIMS IM decreases MS2 spectral complexity in a DDA PASEF workflow.

**Figure 2:**
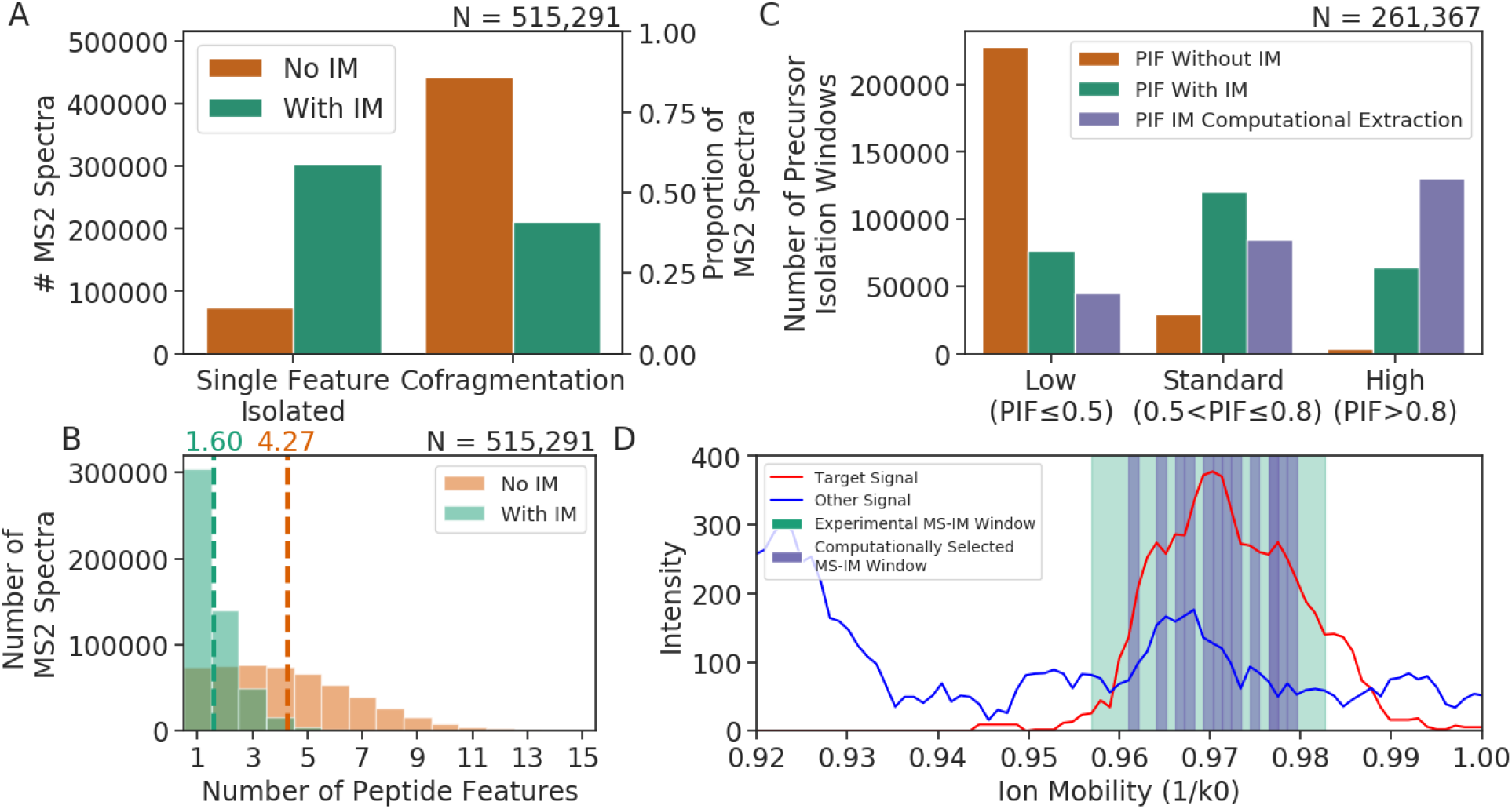
TIMS IM decreases spectral complexity in a DDA-PASEF workflow. (A) Cofragmentation rates in a simulated instrument acquisition with (green) and without (orange) IM. IM separation reduces cofragmentation rates 2.1-fold and increases the number of spectra with a single feature isolated 4.1-fold. (B) MS2 spectra binned by the number of isolated features. Dashed lines indicate the mean of each distribution. IM decreases the average number of features cofragmented per MS2 Spectrum 2.6-fold. (C) Effect of IM separation on precursor isolation window purity. Isolation window quality is assessed by PIF, the proportion of signals that can be attributed to the target precursor. IM separation (green) decreases the number of low quality isolation windows 3.0-fold and increases the number of medium and high quality isolation windows 4.1-fold and 17-fold respectively over non IM isolation windows (orange). Computational extraction of the highest quality IM scans (purple) increases the number of high quality isolation windows 2.0-fold over the non extracted IM window. (D) Ion mobilogram illustrating the computational extraction process. For each isolation window, high quality TOF pushes are selected (purple) based on their computed PIF across a single scan and a new aggregate isolation window is created.

Since timsTOF data acquires 25 TOF pushes per MS2 spectrum which are then aggregated (15 439 300 MS2 TOF pushes total over the course of a 120 minute experiment), we wanted to investigate whether MS2 spectral quality could be increased by filtering out low quality TOF pushes. For each isolation window, we discarded 12 TOF pushes, roughly 50% of the total scans, with the lowest PIF (Figure 2D). We chose to discard roughly 50% of the TOF pushes to balance removing low quality scans and keeping enough scans to maintain a high signal to noise ratio. After low quality TOF pushes were discarded, the PIF across the new aggregate set of TOF pushes was computed. This post-acquisition computation extraction increased the number of high quality isolation windows 2.0-fold (from 24.6% to 50.1%) over the raw experimental data with IM. The highest quality TOF pushes were typically found in the center of the IM isolation window (Supplemental Figure 5), incidentally also providing highest signal to noise. This suggests that by removing low quality TOF pushes, the DDA-PASEF workflow can be optimized post acquisition to further decrease background noise and overall spectral complexity. Since high quality isolation windows are associated with high quality MS2 spectra and higher rates of identification, this post-acquisition computational extraction may allow for deeper proteomic coverage.

The impact of IM on spectral complexity can be further illustrated by comparing a representative MS1 spectrum slice and its corresponding MS2 spectrum: Without IM separation (Figure 3A) only 16% of the ion beam isolated by the quadruple can be attributed to the target precursor. However after IM separation, 67% of signals stem from the target precursor which can be enhanced to 95% of total signals after computational discarding the lowest 12 TOF pushes (Figure 3A). This computational filtering translates to a 34.9% increase (from 0.86 to 1.16) in signal to noise on the corresponding MS2 spectrum and filters out a mislabelled target signal (Figure 3B,C).

**Figure 3:**
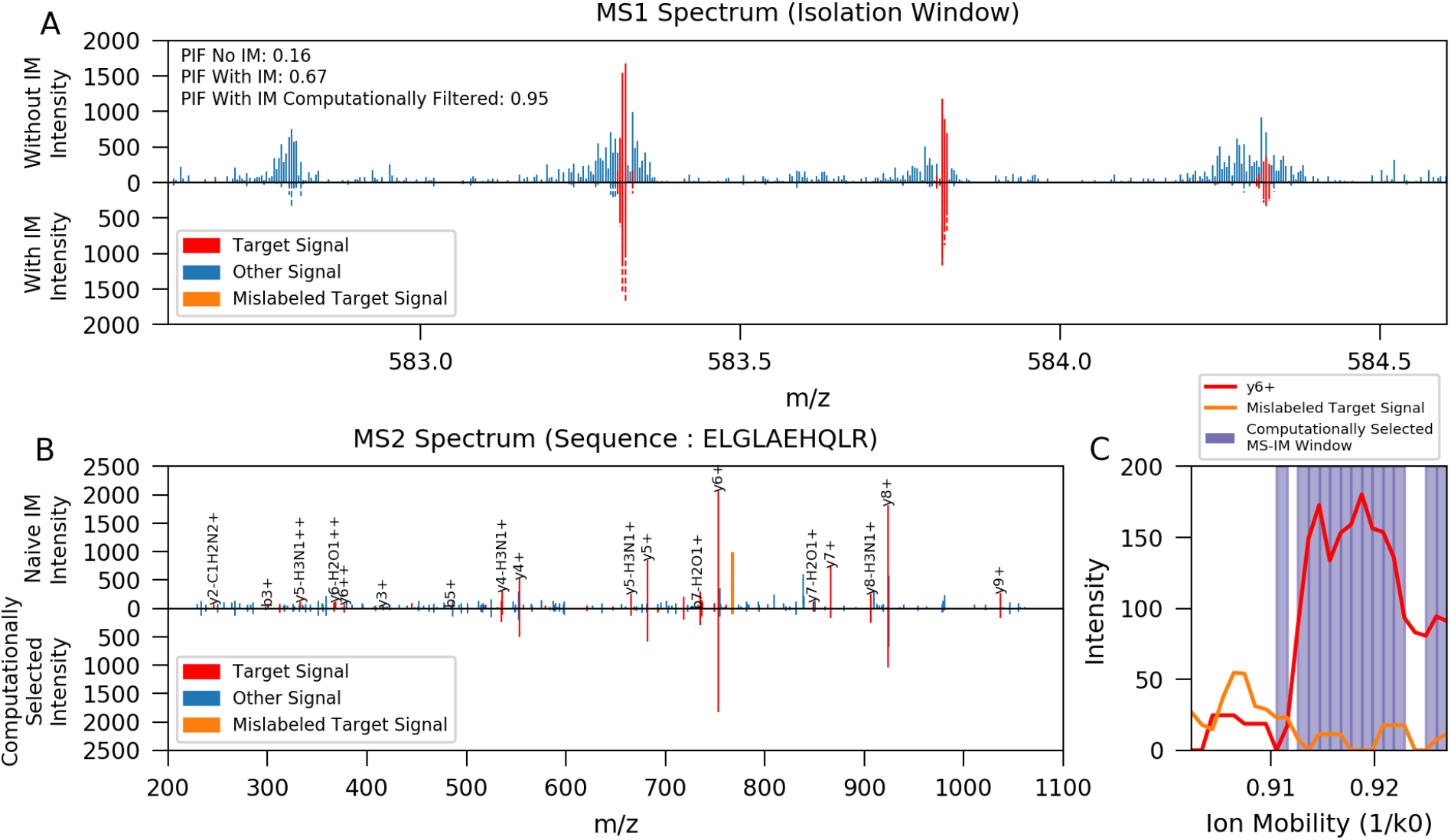
Ion mobility reduces the complexity of spectra. Illustrative example demonstrating how IM reduces background noise while maintaining the target peptide signal. (A) Section of an MS1 spectrum indicative of the instrument chosen isolation window for isolating the target peptide (red). Other signals detected are shown in blue. The top axis shows signals captured without IM and the bottom axis shows signals with IM isolation. On the bottom axis, dashed lines indicate signals in the IM spectrum that do not pass computational filtering, and solid lines indicate IM signals passing the computational filtering. (B) MS2 Fragment ion spectrum resulting from the above MS1 spectrum. The spectrum with IM separation is shown on the top and the spectrum after computational filtering is shown on the bottom. One of the labelled target signals (orange) disappears from the filtered IM spectrum suggesting that it is mislabeled. (C) Ion mobilogram comparing the y6+ and mislabelled ion currents. Computational filtering removes the majority of the mislabelled signal while retaining the y6+ ion current.

### TIMS IM Decreases spectral complexity in diaPASEF-like frames

SWATH-MS is a Data Independent Acquisition (DIA) workflow which cycles through the entire MS2 window in large SWATHs to sample a large portion of the peptide space without bias.^25^ Recently, we developed diaPASEF, a method which combines the SWATH-MS DIA workflow with the PASEF acquisition scheme to sample the IM enhanced peptide space in an unbiased manner. In this workflow, isolation windows have a width of 25 m/z in the MS dimension and an IM width of approximately 0.26 1/k0. Upon data extraction, IM widths are computationally filtered to a size of 0.06 1/k0 to increase sensitivity. ^26^ To quantify the effects of TIMS IM separation in the context of a DIA experiment, we determined the impact of IM on cofragmentation in isolation windows indicative of a DIA workflow. Cofragmentation is an inherent feature in DIA workflows, however the high multiplicity of DIA fragment ion spectra often prevents the unique association of precursor ions with fragment ions (otherwise common in mass spectrometry). Cofragmentation also limits the intra-scan dynamic range of MS2 scans, often leading to limited sensitivity when low intensity peptides are isolated in a DIA window together with high intensity ions. We used the experimental distribution of features derived from a DDA experiment to simulate a DIA experiment with and without IM separation (Figure 4A).

**Figure 4:**
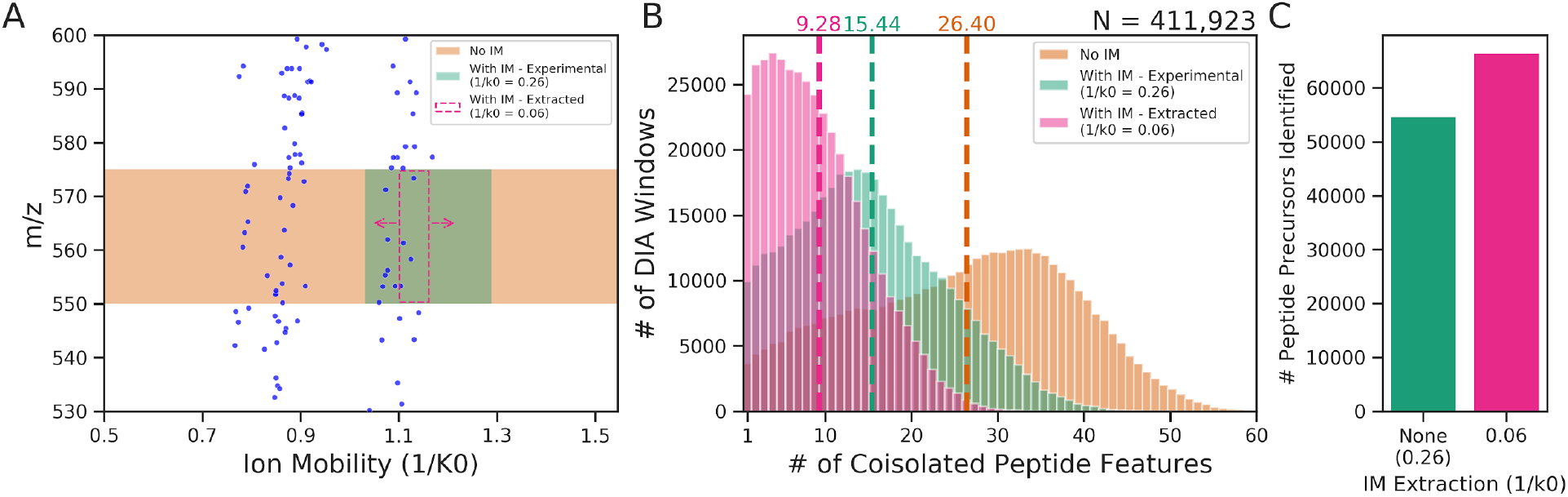
TIMS IM reduces the number of isolated features in simulated DIA windows. (A) Illustration of theoretical isolation windows simulated overlaid on top of peptide feature data. In diaPASEF, MS-IM windows have a fixed location and a width of 0.26 1/k0 (green) and windows are computationally extracted around the precursor of interest to be 0.06 1/k0 wide. (B) Simulated isolation windows with varying IM widths and a m/z width of 25 Da binned by the number of coisolated features. Dashed lines indicate the mean of each distribution. IM isolation widths of 0.06 and 0.26 1/k0 coisolate 2.8-fold and 1.7-fold less features respectively compared to isolation windows without IM. (C) Peptide precursor identification rates 0.06 and 0.26 1/k0 resolutions. An increase in IM selectivity translates to a 21.3% increase in peptide precursor identification rates.

For each theoretical isolation window, we measured the degree of cofragmentation by computing the number of features that overlapped with the isolation window. The addition of IM separation directly decreased the average number of cofragmented peptides from 26.40 to 15.44 (Figure 4B) which is similar in complexity to a 12.5Da m/z isolation window without IM (mean 14.63) (Supplemental Figure 6A). Using the post-acquisition data extraction window of 0.06 1/k0 we found an average of 9.28 cofragmented peptides were isolated (Figure 4B) and this is similar to complexity of a 6.5Da m/z isolation window (mean 8.96) (Supplemental Figure 6B). This observed reduction in the degree of cofragmentation achieved through TIMS IM separation, demonstrates the capability of diaPASEF experiments to cover the majority of the peptide space with high selectivity equivalent to that of 6.5Da m/z isolation windows. The impact of a decrease in spectral complexity, resulting from a decrease in IM width, becomes evident when examining the peptide precursor identification rates in a diaPASEF experiment. Increasing IM separation power through computational extraction (0.26 to 0.06 1/k0 resolution) increases the number of peptide precursors identified by 21.3% (from 54 792 to 66 487) (Figure 4C).^26^ This directly demonstrates how the decrease in spectral complexity resulting from ion mobility translates to an increase in peptide identification rates.

### TIMS IM increases peptide identification rate

The effect of TIMS IM on peptide identification in a DDA-PASEF workflow was evaluated using a categorical model based on previously computed PIF values. We hypothesize that our computed PIF values can be used to estimate peptide identification rates since previous studies suggest that higher PIF values are associated with higher rates of identification.^9^ We also observed this trend in our data, since the median PIF for the identified precursors is greater than the median PIF for targeted and unidentified precursors (Figure 5B). Thus we derived a categorical model by binning precursors by their IM enhanced PIF values and computing the rates of identification of each bin (Figure 5C). By fitting the model to the PIF distribution of non IM enhanced precursors, we estimate that there would be 20 574 identified PSMs in an equivalent experiment without IM, which is 2.05-fold less than the identified PSMs with IM (N = 42 077) (Figure. 5A). Most PSMs without IM separation occur in low quality spectra, defined by their low PIF value, which is in contrast with IM separation PSMs which occur in high quality spectrum (Figure 5D). This number of PSMs without IM is likely an overestimate because it does not correct for the high sequencing speed of PASEF, that is unachievable without TIMS IM separation, and a lower confidence of identification is associated with low PIF values. Thus, consistent with the observations of others,^28^ we argue that IM contributes to at least a 2-fold increase in peptide identification rates.

**Figure 5:**
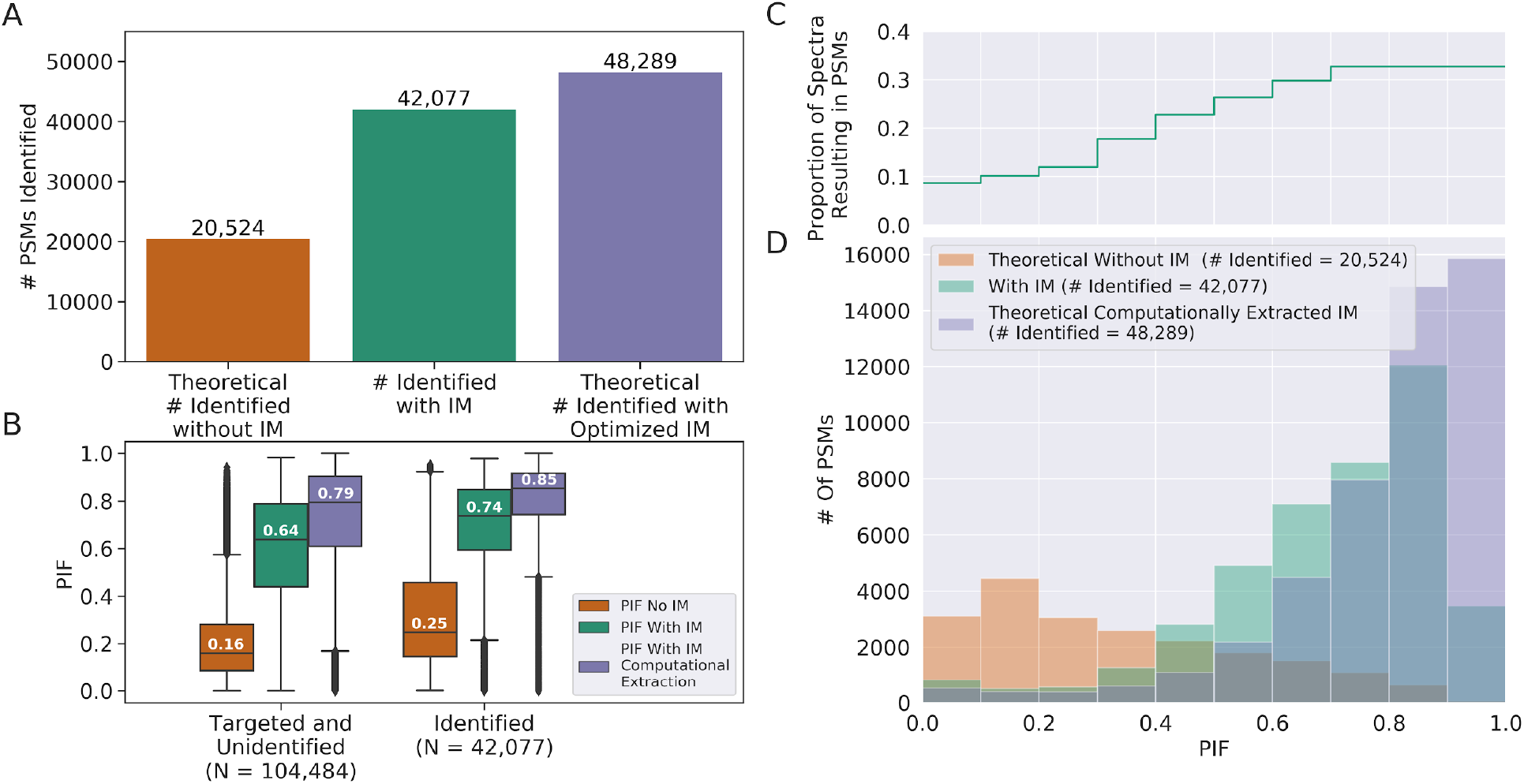
TIMS IM increases peptide identification rates. The PIF, defined as the proportion of signals in an isolation window that can be attributed to the target precursor, was used to estimate the theoretical number of PSMs without IM and with computational extraction along the IM dimension. (A) Comparison of the true number of PSMs in experimental data with IM (green) to the predicted number of PSM without IM (orange) and computational extraction scheme (purple). Model predictions indicate that IM separation leads to an 2.1-fold increase in PSMs. (B) PIF distributions of identified and unidentified precursors. Distributions are also summarized by the median shown. For all distributions, the PIF is greater for identified precursors compared to unidentified precursors, suggesting that identified peptides have a higher PIF than non identified peptides. (C) Illustration of the model used to estimate the number of PSM. This model predicts the proportion of MS2 spectrum leading to PSM as a function of PIF. (D) Distributions of PSMs as a function of PIF with IM (green), without IM (orange) and computationally extracted scheme with IM (purple). Most of the predicted PSMs without IM occur in low quality (PIF ≤ 0.5) spectrum and most of the PSMs with IM and predicted PSMs in the computational extraction scheme occur in standard (0.5 < PIF ≤ 0.8) and high quality (PIF > 0.8) spectrum.

Next, we examined if increasing spectral purity could lead to higher identification rates. We used our model to estimate the number of PSM that would be observed after computational filtering to only high quality IM scans and estimate that filtering leads to a 14.8% increase in the number of PSMs (from 42 077 to 48 289) (Figure 5A). Although computational extraction may not greatly increase identification rates, the observed increase in PIF may result in higher spectral quality and a greater identification confidence.

Computing PIF of untargeted peptides with and without IM shows that IM separation greatly increases the PIF of untargeted features (Supplemental Figure 7). Since many untargeted peptides undergoing IM separation have a high PIF, we hypothesise that an increase in sequencing speed would greatly increase the number of PSM with IM separation and would not have as large of an effect on increasing the number of PSM without IM. To test this hypothesis, we applied the model above to estimate the additional number of PSMs that would result by targeting all untargeted peptides with and without IM selection (Supplemental Figure 8B). We estimate that if all untargeted features were targeted, there would be an additional 43 950 PSM with IM in contrast to 24 213 additional PSMs without IM (Supplemental Figure 8A). Thus, similar to above, the PSM with IM are likely of higher confidence due to their greater PIF values of their identifications (Supplemental Figure 8C) meaning that any increase in sequencing speed would disproportionately benefit the IM-enabled workflow.

## Discussion

As MS sequencing speed has been rising over the last instrument generations, one of the increasingly more appreciated impediments to high proteome coverage in shotgun proteomics workflows is high spectral complexity. ^29^ In complex samples, the large number of analytes concurrently being ionized prevents the isolation of pure analytes using a quadrupole mass filter, which impacts identification and quantification.^1,3,5^ Increased on-line or off-line peptide separation has been used with great success to decrease cofragmentation rates and boost identification and quantification performance. One such peptide separation strategy is IM, which separates peptides by their collisional cross section on a sub millisecond timescale. TIMS IM has been integrated into the PASEF workflow resulting in high protein and peptide identification rates in complex proteomic samples.^24,26^ However, there are multiple factors that may explain the benefits of this workflow including; ion trapping, increased sequencing speed, ion focusing and IM separation. Since all of these effects are achieved through use of the TIMS device, it is difficult to distinguish their contributions on improving peptide identification. To understand the mechanisms responsible for improved peptide identification observed in the PASEF workflow, it is necessary to study each of the above factors individually and in isolation of one another. In this study, we quantified the impact of one of these factors, IM separation, by modelling experimental PASEF data from a HeLa cell digest with and without IM separation to quantify the IM separation power of TIMS.

Coupling TIMS IM separation to liquid chromatography and a mass analyzer adds an additional, partially orthogonal, dimension of separation to the mass spectrometric workflow. This additional separation results in the resolution of interfering peptides. To quantify the extent that TIMS IM separation reduces peptide interference, we modelled peptide features with and without IM separation and computed the global pairwise overlaps. We found that IM increases the proportion of peptides without overlap 9.2 fold, consistent with the TIMS separation power observed by Meier et al. ^28^ Although IM peak capacity may be lower than retention time or m/z peak capacity (Supplemental Figure 3), coupling IM to the mass spectrometric workflow provides substantial reduction to peptide interferences in complex peptide samples. Delving deeper into the nature of the species being separated by TIMS IM we found that the majority of the separation power results from separation of peptides from each other rather than peptides from contaminants (Supplementary Results). These findings suggest that TIMS IM is not merely separating out contaminants, but also provides substantial simplification to the overall peptide space.

To evaluate the benefits of IM separation in the context of a DDA-PASEF workflow, we computed the impact of IM separation on spectral complexity and peptide identification rates. Isolation windows were derived from a DDA-PASEF experiment limiting assumptions required on the instrument acquisition cycle. MS2 spectral complexity was inferred from the MS1 level by estimating cofragmentation rates and spectral quality from the isolation events. We found that TIMS IM separation decreases the rates of cofragmentatation 2.1-fold and increases the number of spectra without cofragmentation 4.1 fold. These results are consistent with the 3.0-fold reduction in low quality isolation windows and 4.1-fold improvement in standard quality isolation windows observed with the addition of IM separation. Moreover, we show that low quality TOF pushes can be discarded to increase the number of high quality scans 2.0-fold while maintaining a high signal-to-noise ratio. Using the computed PIF values, we estimated that IM improves the rates of PSM at least 2.0-fold, since our model does not take into account the increased sequencing speed permissible with TIMS IM separation. The reduction in spectral complexity and the estimated improvement in PSMs resulting from TIMS IM separation, demonstrate the benefits of incorporating TIMS IM separation to improve peptide identification rates in DDA experiments.

DIA isolation windows are widened to enable sufficient sampling of the entire peptide space peptide however, this results in complex fragment ion spectra with a high degree of cofragmentation. We show that by coupling DIA to IM, the spectral complexity is greatly reduced. Using simulated DIA isolation windows indicative of a diaPASEF experiment, we found that the standard IM isolation window used in a diaPASEF experiment reduces the mean number of peptides isolated per spectrum 2.8-fold, which we estimate has similar complexity to a 6.5Da isolation window width without IM. Furthermore, as IM measurements increase in accuracy, smaller DIA extraction windows can be used which we expect to linearly decrease the number of cofragmented peptides resulting in increased peptide identification rates. (Supplemental Results). These findings suggest that IM separation has the potential to provide major benefits to DIA workflows by reducing the number of peptides isolated per window without sacrificing sequencing depth.

It is important to consider that our quantifications of TIMS IM separation are limited by the accuracy of the MaxQuant feature detection algorithm since it assumes that all MaxQuant detecting features are true features. Feature detection algorithms are well established for detecting features across the RT and m/z dimensions however, due to the novelty of widespread use to IM-MS, peptide feature detection across the RT, m/z and IM dimensions may not have the same accuracy. Increased separation may lead to the detection of more noise peaks which may be assembled into false peptide features, causing an overestimation of the number of peptide features present and consequently resulting in an overestimation of the separation power of TIMS IM. Although these assumptions may limit the quantification accuracy of TIMS IM separation power, we believe the observed effect is sufficiently large and accurate to highlight the impact of TIMS IM separation in a bottom up proteomics workflow.

We conclude that separation by TIMS IM in the context of PASEF acquisition increases overall separation of analytes and provides significant benefits to DDA and DIA proteomic workflows by decreasing spectral complexity. This suggests that IM separation contributes considerably to the high protein identification rates observed in a PASEF acquisition workflow. Furthermore, the observed decrease in cofragmentation rates is consistent with the findings that TIMS IM separation improves MS2 label based quantification techniques.^18^ These findings highlight the benefit of IM separation in bottom up proteomic experiments of complex samples in achieving high peptide identification and accurate peptide quantification. Due to the novelty of TIMS device, we also expect future iterations of the device to have additional capabilities and resolving power, which could further improve upon identification and quantification accuracy.

## Methods

### Data Acquisition and Feature Extraction

Simulations and computations were conducted on data published by Meier et al.^24^ This data was acquired from a HeLa cell digest acquired on an online trapped ion mobility spectrometry - quadrupole time of flight mass spectrometer (timsTOF Pro, Bruker Daltonics) in PASEF acquisition mode (PXD010012). Analysis was performed on the second technical replicate of the 100ms TIMS accumulation time. The 100ms TIMS accumulation time was used because it was previously established as optimal.^24^ Features were previously computed by Meier et al. using the MaxQuant framework.^27^

### Computational Framework

Pairwise feature overlap and overlap between isolation windows and peptide features were computed in Python with the assistance of the NCLS (v0.0.53) ^30^ and numpy (v1.19.1)^31^ packages. Data analysis was conducted in the JupyterLab Framework and manipulation was conducted with the Pandas (v1.1.3) ^32,33^ package. Plots were generated using the matplotlib (v3.1.3)^34^ and seaborn(v0.11.0)^35^ libraries. Source code for analyses below can be found at https://github.com/jcharkow/imMQExplorer.

### Assessing the effects of IM on Feature Density

To assess the impact of IM separation on peptide density, peptide features detected by MaxQuant were modelled with and without the IM dimension. In this model, a peptide feature is defined as an isotopic envelope across retention time (RT) and IM. The width of the isotopic envelope was defined based on the m/z, charge and number of isotopic peaks reported by MaxQuant. The elution time of the feature was defined by the RT and the retention full width half maximum (FWHM) length reported by MaxQuant. The IM profile of the feature was defined by the IM index and the IM index FWHM length. To compute the effect of IM on feature density, all pairwise feature overlaps were computed with and without the IM dimension. Overlap in the RT and IM dimensions were defined as features with overlap in their FWHM elution profiles. Overlap in m/z was defined as overlap in the features’ isotopic envelopes, as such features could not be independently isolated by a standard quadrupole selection, and are thus likely cofragmented.

To evaluate the relationship between IM separation power and feature density, IM separation power was computed across m/z and RT dimensions as feature density varies throughout these dimensions. To compute IM separation power, the sliding mean of the proportion of features with greater than a set number of overlaps was computed with and without IM and these computations. The relationship between IM separation power and feature density was determined by comparing the above functions in high and low feature density regions.

### Assessing the effects of IM on peptide cofragmentation in the context of a DDA experiment

To measure the effect of IM on spectral complexity, the rates of peptide cofragmentation with and without IM separation were computed. Two sets of isolation windows were constructed, one with IM isolation and the other set without IM. Isolation windows with IM were derived from experimental instrument acquisition data from a TimsTOF pro in DDA-PASEF mode. Isolation windows without IM were constructed by removing the IM dimension from the IM isolation windows. The degree of cofragmentation was measured by computing the number of features overlapping with each isolation window. Overlap in the RT and IM space is defined as overlap in feature FWHM with the isolation window’s IM isolation range or point in RT respectively. M/z overlap is defined as overlap of the feature’s isotopic envelope with the quadrupole m/z isolation window. This assumes an isolation window overlapping with two or more isotopic traces of a peptide will lead to interfering fragment ions. Since the instrument acquisition did not always target precursors in their RT or IM FWHM region, this resulted in some isolation windows reporting isolation of no features. These pairs of isolation windows were excluded from the analysis (N = 102 281 in each set) resulting in a total of 515 291 isolation windows in each set. To compute the rates of cofragmentation each isolation window was classified as either containing or not containing a cofragmentation event. A cofragmentation event was defined as an isolation window overlapping with more than one peptide feature. The degree of cofragmentation was assessed by determining the number of features isolated per window.

### Evaluating the effects of IM on spectral complexity using Precursor Ion Fraction (PIF)

The precursor ion fraction (PIF) is measurement on the MS1 level data defined as the proportion of the isolated ion beam that stems from targeted signals. For each targeted feature, the MS1 spectrum which triggered the isolation of the feature was determined using the instrument acquisition data. The MS1 spectrum was filtered to only contain the signal within the instrument’s reported isolation parameters in m/z and IM. For PIF computations without IM, filtering by IM isolation was omitted. The PIF is computed as the proportion of intensity in the filtered MS1 spectrum that can be attributed to the target feature. Peaks were assigned to the target feature if they were within 20 ppm of any of the isotopic traces of the analyte and also matched the target feature position in RT and IM. Note that this approach likely overestimates PIF slightly since interfering signals closer than 20 ppm to the target feature would be attributed to the target feature, making our analysis err on the side of being more conservative.

To compute the PIF for untargeted features, features were mapped to the closest MS1 spectrum that occurred before its MaxQuant reported retention time. The MS1 spectrum was filtered to 2 Da and 25 ion mobility scans (0.026 1/k0) wide centered around the feature, typical of an average isolation window for this instrument. To compute PIF without IM, the IM filtering was omitted.

### Computational Extraction of High Quality TOF Pushes

Computational extraction along the IM dimension can occur because each IM isolation window is composed of a discrete set of 25 TOF pushes, with the full isolation window formed of an aggregate of all these pushes. Since each TOF push is easily separable from one another, each TOF push can be analyzed individually and a subset of the pushes can be merged to create a new spectrum. To filter out low quality TOF pushes, the PIF was computed for each individual TOF push and the 13 pushes with the highest PIF were extracted and merged to create a new spectrum. The aggregate was composed of 13 pushes since this corresponds to roughly 50% of the TOF pushes and excluding too many pushes may result in a decreased signal-to-noise ratio. The PIF was then computed on the aggregate spectrum as described above.

### Effect of IM on the degree of cofragmentation in a DIA workflow

To evaluate the effects of IM in a DIA workflow, theoretical isolation windows were constructed based on MaxQuant feature characteristics to be representative of a diaPASEF experiment. The degree of coisolation was determined by computing the number of features which overlapped with each isolation window. Multiple sets of theoretical isolation windows were constructed for each feature, varying by their m/z and IM isolation width, and uncentered around the feature. Isolation windows were either 50, 25, 12 or 6.5 Da wide in the m/z dimension (see supplemental analyses for 50, 12 and 6.5Da windows). In the IM dimension, various IM widths were constructed illustrative of diaPASEF experimental (around 0.25 1/k0) width, computationally extracted (around 0.06 1/k0) width, or spanning the entire IM dimension to simulate no IM separation being present. Overlap between isolation windows and features were computed as described for DDA isolation windows.

To determine the effects of IM on peptide identification rate, OpenSwath and Pyprophet were run as described in Meier et al.^26^ on the 200ng diaPASEF runs (PXD017703). The only change from the described methods was that the ION_MOBILITY_WINDOW parameter was set to 0.06 and -1 to evaluate the impact of a reduced IM resolution.

### Estimating the number of PSM using PIF values

To evaluate the influence IM has on peptide identification, we predicted the expected number of PSMs without IM and compared this to the number of PSMs with IM. To accomplish this, we created and applied a categorical model which estimates the likelihood a targeted peptide will be identified based on the precursor’s PIF value. To derive the model, each targeted precursor from a DDA PASEF experiment was binned by its computed PIF value (bin width 0.1). For each bin, we computed the proportion of targeted precursors that resulted in a PSM. The bin between 0.9-1.0 was set to the same proportion as the 0.8-0.9 bin because of an unexpected drop in proportion of PSMs, likely due to a small amount of data points in this bin.

To estimate the number of PSMs that would occur in an equivalent experiment without IM, PIF values without IM were binned and the number of features per bin was multiplied by the model’s proportion of targeted features that resulted in a PSM. The model was also applied to the PIF values computed after computational extraction along the IM dimension.

## Supporting information

Supplemental Results

## Supporting Information

1. Supplemental Figures including: pairwise overlap comparison with and without IM, relationship between IM and feature density, separation power of RT, m/z and IM, distribution of PIF, histogram of highest quality scans, comparison of MS and MS-IM theoretical DIA isolation windows, PIF of untargeted peptide features and predicted identification rates of untargeted peptides. (PDF)
2. Comparison of background-peptide and peptide-peptide separation power, comparison of intercharge and intracharge separation power and, relationship between IM window size and spectra complexity. (PDF)

## Acknowledgements

The authors would like to thank Annie Ha, Aparna Srinivasan and Tom Oulette for their contributions to the project. This Project was funded by the Government of Canada through Genome Canada and the Ontario Genomics Institute (OGI-164). J. C. was supported by the Ontario Graduate Scholarship. H.L.R. is supported by the Canadian Foundation for Innovation and the John R. Evans Leaders Fund and is the Canada Research Chair in Mass Spectrometry-based Personalized Medicine.

## References

1. Houel, S. et al. Quantifying the Impact of Chimera MS/MS Spectra on Peptide Identification in Large-Scale Proteomics Studies. J. Proteome Res. 9, 4152–4160 (2010).

2. Ow, S. Y., Salim, M., Noirel, J., Evans, C. & Wright, P. C. Minimising iTRAQ ratio compression through understanding LC-MS elution dependence and high-resolution HILIC fractionation. PROTEOMICS 11, 2341–2346 (2011).

3. Bantscheff, M. et al. Robust and sensitive iTRAQ quantification on an LTQ Orbitrap mass spectrometer. Mol. Cell. Proteomics MCP 7, 1702–1713 (2008).

4. Ow, S. Y. et al. iTRAQ Underestimation in Simple and Complex Mixtures: “The Good, the Bad and the Ugly”. J. Proteome Res. 8, 5347–5355 (2009).

5. DeSouza, L. V. et al. Multiple Reaction Monitoring of mTRAQ-Labeled Peptides Enables Absolute Quantification of Endogenous Levels of a Potential Cancer Marker in Cancerous and Normal Endometrial Tissues. J. Proteome Res. 7, 3525–3534 (2008).

6. Li, H. et al. Estimating Influence of Cofragmentation on Peptide Quantification and Identification in iTRAQ Experiments by Simulating Multiplexed Spectra. J. Proteome Res. 13, 3488–3497 (2014).

7. Cox, J. et al. Andromeda: A Peptide Search Engine Integrated into the MaxQuant Environment. J. Proteome Res. 10, 1794–1805 (2011).

8. Zhang, B., Pirmoradian, M., Chernobrovkin, A. & Zubarev, R. A. DeMix Workflow for Efficient Identification of Cofragmented Peptides in High Resolution Data-dependent Tandem Mass Spectrometry. Mol. Cell. Proteomics 13, 3211–3223 (2014).

9. Michalski, A., Cox, J. & Mann, M. More than 100,000 Detectable Peptide Species Elute in Single Shotgun Proteomics Runs but the Majority is Inaccessible to Data-Dependent LC−MS/MS. J. Proteome Res. 10, 1785–1793 (2011).

10. Ting, L., Rad, R., Gygi, S. P. & Haas, W. MS3 eliminates ratio distortion in isobaric multiplexed quantitative proteomics. Nat. Methods 8, 937–940 (2011).

11. Canterbury, J. D., Yi, X., Hoopmann, M. R. & MacCoss, M. J. Assessing the dynamic range and peak capacity of nanoflow LC-FAIMS-MS on an ion trap mass spectrometer for proteomics. Anal. Chem. 80, 6888 (2008).

12. Baker, E. S. et al. An LC-IMS-MS Platform Providing Increased Dynamic Range for High-Throughput Proteomic Studies. J. Proteome Res. 9, 997 (2010).

13. Saba, J., Bonneil, E., Pomiès, C., Eng, K. & Thibault, P. Enhanced Sensitivity in Proteomics Experiments Using FAIMS Coupled with a Hybrid Linear Ion Trap/Orbitrap Mass Spectrometer †. J. Proteome Res. 8, 3355–3366 (2009).

14. Venne, K., Bonneil, E., Eng, K. & Thibault, P. Improvement in Peptide Detection for Proteomics Analyses Using NanoLC−MS and High-Field Asymmetry Waveform Ion Mobility Mass Spectrometry. Anal. Chem. 77, 2176–2186 (2005).

15. Pfammatter, S., Bonneil, E. & Thibault, P. Improvement of Quantitative Measurements in Multiplex Proteomics Using High-Field Asymmetric Waveform Spectrometry. J. Proteome Res. 15, 4653–4665 (2016).

16. Schweppe, D. K. et al. Characterization and Optimization of Multiplexed Quantitative Analyses Using High-Field Asymmetric-Waveform Ion Mobility Mass Spectrometry. Anal. Chem. 91, 4010–4016 (2019).

17. Baker, E. S. et al. Enhancing bottom-up and top-down proteomic measurements with ion mobility separations. Proteomics 15, 2766–2776 (2015).

18. Ogata, K. & Ishihama, Y. Extending the Separation Space with Trapped Ion Mobility Spectrometry Improves the Accuracy of Isobaric Tag-Based Quantitation in Proteomic LC/MS/MS. Anal. Chem. 92, 8037–8040 (2020).

19. Meier, F. et al. Parallel Accumulation–Serial Fragmentation (PASEF): Multiplying Sequencing Speed and Sensitivity by Synchronized Scans in a Trapped Ion Mobility Device. J. Proteome Res. 14, 5378–5387 (2015).

20. Cumeras, R., Figueras, E., Davis, C. E., Baumbach, J. I. & Gràcia, I. Review on Ion Mobility Spectrometry. Part 1: current instrumentation. The Analyst 140, 1376–1390 (2015).

21. Fernandez-Lima, F. A., Kaplan, D. A. & Park, M. A. Note: Integration of trapped ion mobility spectrometry with mass spectrometry. Rev. Sci. Instrum. 82, 126106 (2011).

22. Fernandez-Lima, F., Kaplan, D. A., Suetering, J. & Park, M. A. Gas-phase separation using a trapped ion mobility spectrometer. Int. J. Ion Mobil. Spectrom. 14, 93–98 (2011).

23. Silveira, J. A., Ridgeway, M. E., Laukien, F. H., Mann, M. & Park, M. A. Parallel accumulation for 100% duty cycle trapped ion mobility-mass spectrometry. Int. J. Mass Spectrom. 413, 168–175 (2017).

24. Meier, F. et al. Online parallel accumulation – serial fragmentation (PASEF) with a novel trapped ion mobility mass spectrometer. Mol. Cell. Proteomics (2018) doi:10.1074/mcp.TIR118.000900.

25. Gillet, L. C. et al. Targeted Data Extraction of the MS/MS Spectra Generated by Data-independent Acquisition: A New Concept for Consistent and Accurate Proteome Analysis. Mol. Cell. Proteomics MCP 11, (2012).

26. Meier, F. et al. diaPASEF: parallel accumulation–serial fragmentation combined with data-independent acquisition. Nat. Methods 17, 1229–1236 (2020).

27. Prianichnikov, N. et al. MaxQuant Software for Ion Mobility Enhanced Shotgun Proteomics *. Mol. Cell. Proteomics 19, 1058–1069 (2020).

28. Meier, F. et al. Deep learning the collisional cross sections of the peptide universe from a million experimental values. Nat. Commun. 12, 1185 (2021).

29. Shishkova, E., Hebert, A. S. & Coon, J. J. Now, More Than Ever, Proteomics Needs Better Chromatography. Cell Syst. 3, 321–324 (2016).

30. Stovner, E. B. & Sætrom, P. PyRanges: efficient comparison of genomic intervals in Python. Bioinformatics 36, 918–919 (2020).

31. Harris, C. R. et al. Array programming with NumPy. Nature 585, 357–362 (2020).

32. The pandas development team. pandas-dev/pandas: Pandas. (Zenodo, 2020). doi:10.5281/zenodo.3509134.

33. McKinney, W. Data Structures for Statistical Computing in Python. in Proceedings of the 9th Python in Science Conference (eds. Walt S. van der & Millman, J.) 56–61 (2010). doi:10.25080/Majora-92bf1922-00a.

34. Hunter, J. D. Matplotlib: A 2D Graphics Environment. Comput. Sci. Eng. 9, 90–95 (2007).

35. Waskom, M. & the seaborn development team. mwaskom/seaborn. (Zenodo, 2020). doi:10.5281/zenodo.592845.

