## Supplemental Figures for "Trapped Ion Mobility Spectrometry Reduces Spectral Complexity in Mass Spectrometry Based Workflow"

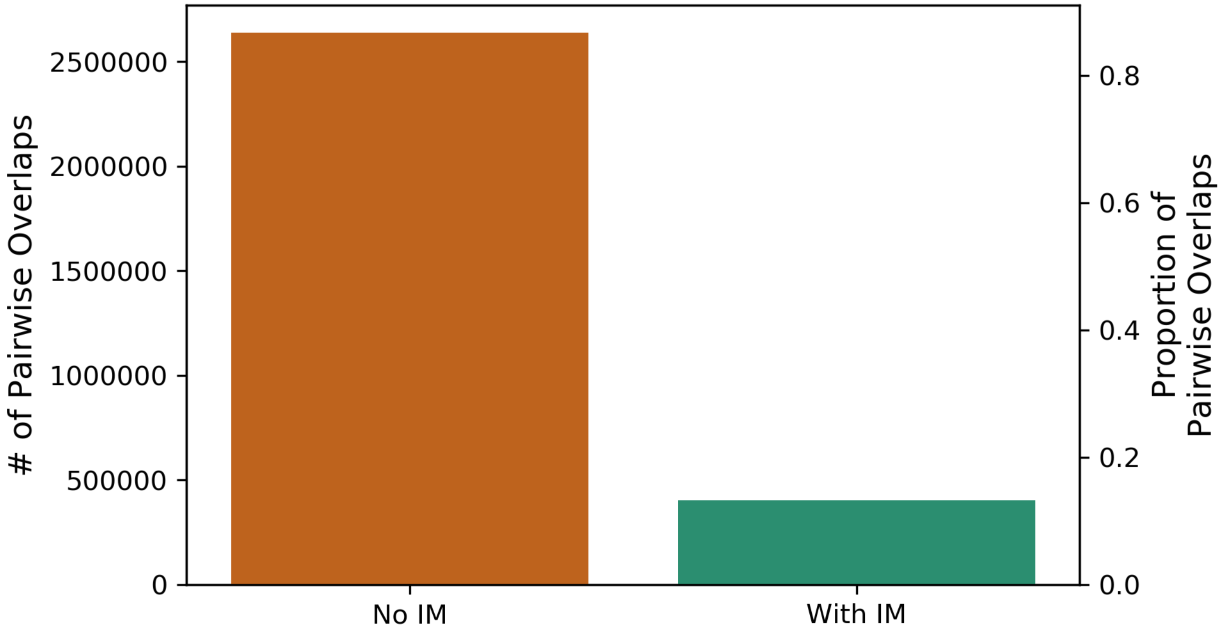


***Supplementary Figure 1: IM reduces peptide feature overlap.*** *Number and proportion of pairwise overlaps between MaxQuant detected peptide features without (orange) and with (green) IM separation. IM reduces the total pairwise overlaps 6.5-fold.*

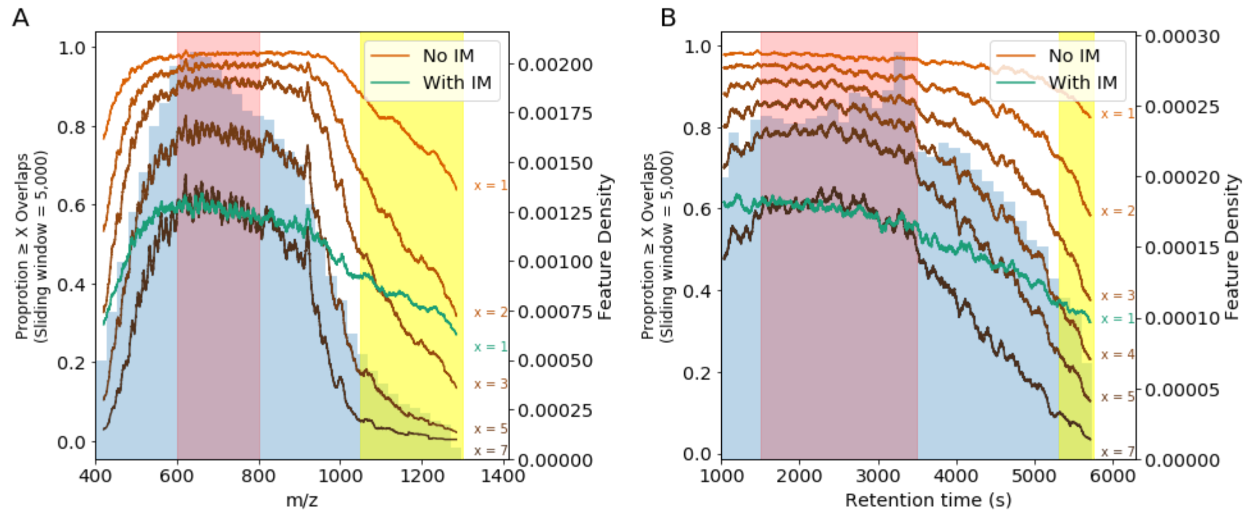


***Supplemental Figure 2: IM separation power is dependent on feature density.*** *Sliding average of the proportion of features with greater than X overlaps without (orange) and with IM (green) throughout the (A) m/z and (B) RT dimensions. These results are overlaid on top of the feature distribution. In the high density regions (red), the proportion of IM features with overlap is similar to the proportion of non IM enhanced features with 7 overlaps suggesting that IM reduces feature density overlap 7-fold in these regions. In low feature density regions (yellow) the proportion of IM features with overlap is similar to the proportion of no-IM features with 3-4 overlaps suggesting that IM reduces feature density approximately 3-4 fold in these regions. This suggests that IM separation power is approximately 2-fold greater in high density regions compared to low density regions.*

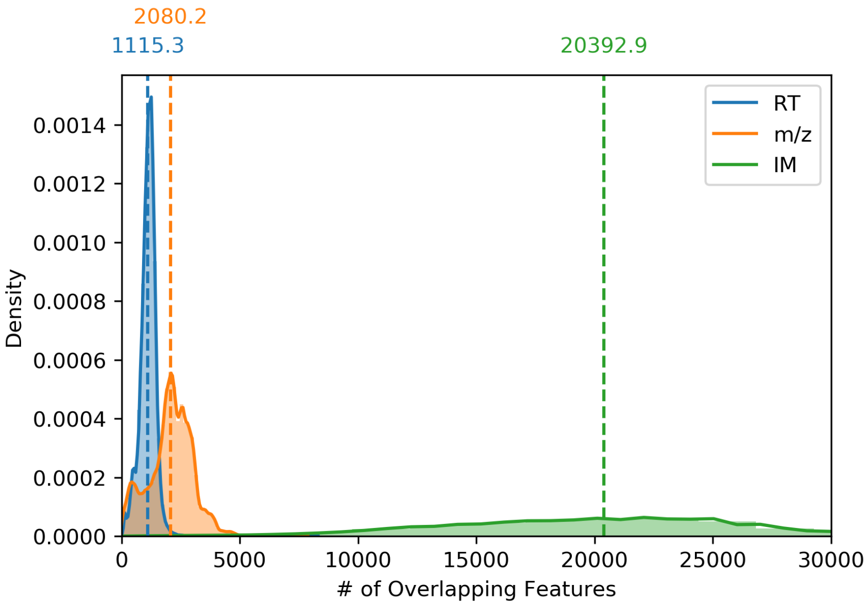


***Supplemental Figure 3 - IM has a lower peak capacity compared to RT and m/z.*** *Kernel density estimation of the number of overlaps per feature for RT, m/z and IM axes individually. Dashed lines represent the mean of each distribution. The average number of overlapping features per peptide across the IM dimension is 18.3-fold and 9.8-fold greater than the number of overlapping features across the RT or m/z dimensions respectively.*

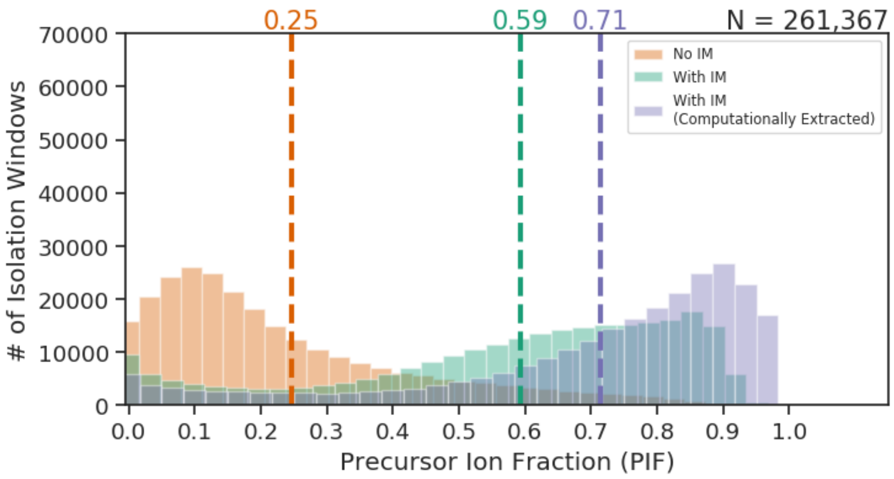


***Supplementary Figure 4: IM increases the PIF of isolation windows.*** *Distribution of isolation windows bined by their Precursor Ion Fraction (PIF) without IM (orange), with IM (green) and computational extracted IM (purple).*

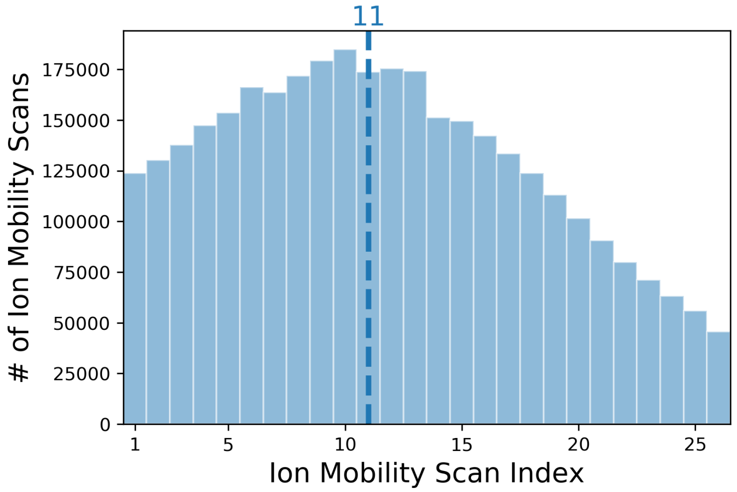


***Supplementary Figure 5: The highest quality IM scans are most commonly found in the center of the IM window.*** *Distribution of the position of the 13 highest quality IM scans per isolation window as evaluated by PIF. The median of the distribution is shown by the dashed blue line.*

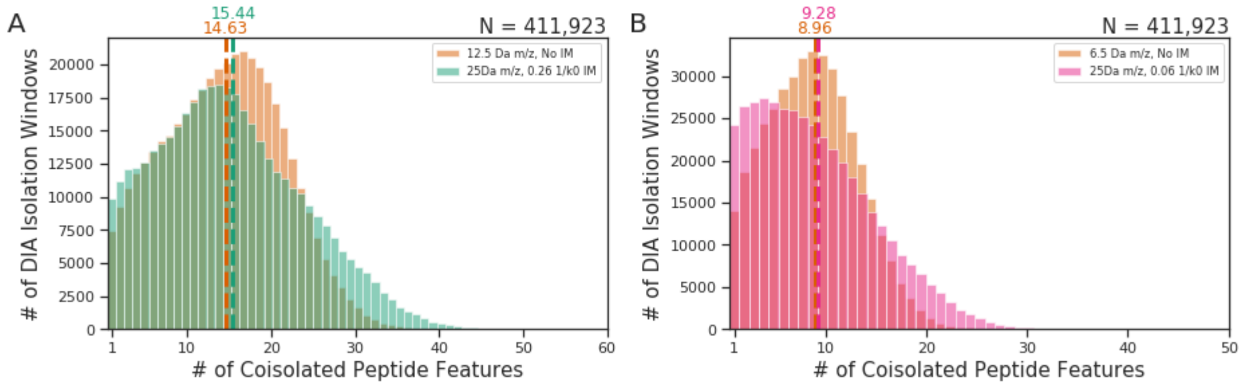


***Supplementary Figure 6: High selectivity MS-IM isolation windows have similar complexity to 6.5Da isolation windows.*** *Comparison of MS-IM and MS isolation coisolation rate distributions. (A) MS-IM windows with 25Da and 0.26 1/k0 have similar coisolation rates as MS windows of 12.5Da. (B) MS-IM windows with 25Da and 0.26 1/k0 has a similar coisolation rates to MS windows of 12.5Da. Means of the distributions are shown as dashed lines.*

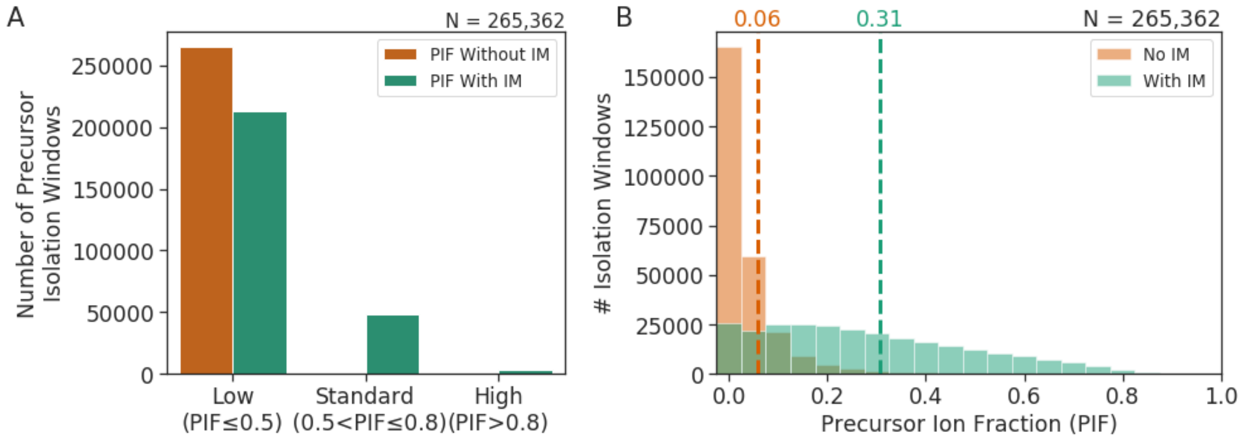


***Supplementary Figure 7: IM increases PIF of untargeted peptides.*** *(A) Effect of IM on isolation window quality for untargeted precursors. IM increases the number of standard and high quality isolation windows of untargeted precursors. (B) Distribution of theoretical IM-enhanced and non-IM enhanced isolation windows constructed around untargeted peptide features binned by their PIF. Dashed lines indicate the mean of each distribution. IM increases the PIF of the average isolation window 5.2 fold.*

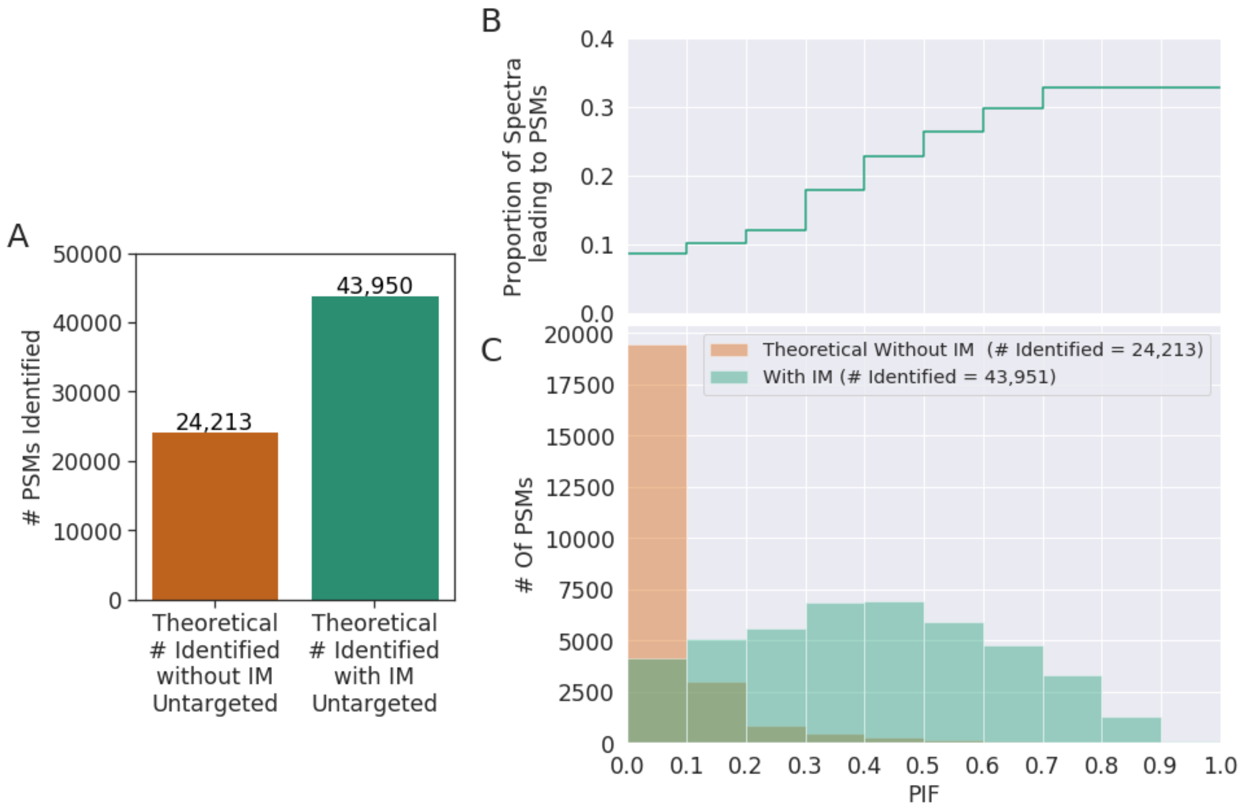


***Supplemental Figure 8: Increasing PASEF sequencing speed disproportionately benefits experiments with IM separation compared to non-IM enhanced data.*** *The increase in PSMs that would result by sequencing untargeted peptides is estimated with and without IM based on the model used in Figure 4. (A) Estimated number of additional PSMs that would result from sequencing untargeted peptides without (orange) and with (green) IM separation. It is estimated that IM would lead 2.1-fold more PSM than non IM enhanced data if all untargeted precursors were targeted. (B) Model used to estimate the proportion of precursors with PSM, derived from the PIF and rate of identification of experimental DDA PASEF targeted peptides. (C) Estimation of the number of PSM of untargeted features with (green) and without (orange) IM. Estimated PSMs without IM occur predominantly in a lower PIF range compared to PSMs with IM.*
