## Supplemental Results for "Trapped Ion Mobility Spectrometry Reduces Spectral Complexity in Mass Spectrometry Based Workflow"

**Peptide-Peptide is more prevalent than Peptide-Background separation**

Since contaminants have greater variation in their molecular properties than peptides do from each other, we expected that the majority of separation power gained from IM would result from the separation of peptides from contaminants. To test this, we classified each MaxQuant identified feature as either a contaminant or a peptide feature based on its charge state. Features with a charge state of 1 were classified as contaminants and features with other charge states were assumed to be true peptide features. For each pairwise overlap between features without IM, we classified the overlaps as resolved by IM occurring between a peptide and a contaminant, resolved by IM occurring between two peptides or unresolved by IM. Features with no overlaps without IM were excluded from the analysis (N = 17 144). For each peptide feature, we computed the proportion of overlaps in each classification group. Surprisingly, we found that for the average peptide feature, the majority of resolved overlaps were between two peptide features rather than between a peptide feature and a contaminant (Figure 1A) and, these trends were consistent across different charge states (Figure 1B-E). This suggests that IM separation power between peptide features is stronger than the IM separation power between peptide features and contaminants.

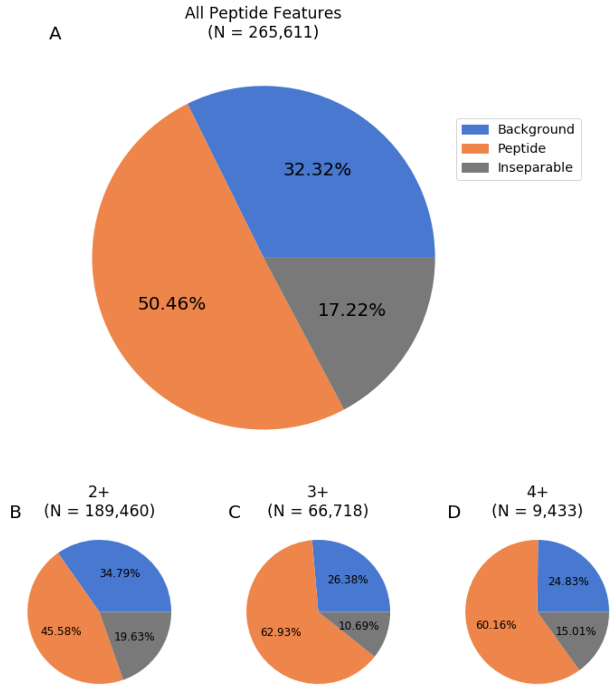


***Figure 1: Peptide-Peptide separation power is stronger than Peptide-Contaminant separation power.*** *Proportion of separations resolved by IM separating peptides (orange), resolved by IM separating peptides from contaminants (blue) and unresolved by IM (grey). The relative separation power is shown for (A) the average peptide feature and for specific charge states (B-D).*

**TIMS IM has a greater impact on intercharge compared to intracharge separation**

An analyte's measured IM is dependent on its collisional cross section, which is reflected both in its m/z and charge value (Figure 2). Due to these dependencies, IM separates peptides both within and between charge states. To investigate the relationship between separation power and charge state, we compared the relative influence of intercharge and intracharge separation power at resolving pairwise feature overlaps. Due to the distribution of peptide features by IM (Figure 2) we expected that charge state plays a major role in separating features. To quantify the relative separation power of intercharge and intracharge separation, for each feature we computed the proportion of total overlaps that were resolved by each separation type. Features with no overlaps without IM were excluded (N= 20 739). In the average feature, about half of the overlaps were resolved by intercharge influences, a third were resolved by intracharge influences and the remaining overlaps were inseparable (Figure 3A). These findings suggest that for the average peptide feature, TIMS IM intercharge separation power is 1.8-fold stronger than intracharge separation power.

Next we considered the intercharge and intracharge influences in each charge state individually. For each charge state, we found that in the average feature, the proportion of overlaps separable by intercharge effects was larger than intracharge effects, however the relative separation power between charge states varied (Figure 3B-E). The estimated intracharge separation power was 1.5 fold larger in 2+ charge state compared to the 3+ charge state. (Figure 3C, D). This trend may be observed because the 2+ species make up a greater proportion of the total features compared to 3+ species. Therefore, separating charge states removes more interfering ions from the 3+ species than from the 2+ species.  A charge state group with a greater number of analytes corresponds with a larger proportion of analytes that cannot be separated by charge state and thus must be more dependent on intracharge separation. This interpretation is consistent with the trends observed that as the proportion of total features contained in a charge state increases, interchange influence decreases and intracharge influence increases (Figure 4). Although for all charge states, intercharge separation power is greater than intracharge separation power, the intercharge separation power is found to be weaker in more populous charge state groups.

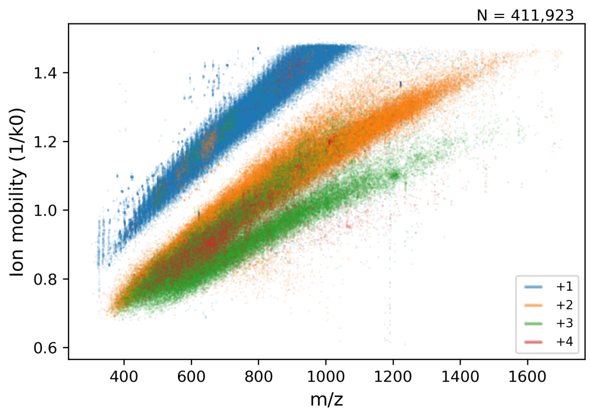


***Figure 2 - Relationship between IM, m/z and charge state for features from a DDA-PASEF experiment.*** *Scatterplot of peptide features monoisotopic m/z by IM, coloured by charge state in a DDA PASEF Experiment. Features of the same charge state form distinct plumes. This figure was redrawn based on data from Meier et al.*[1]

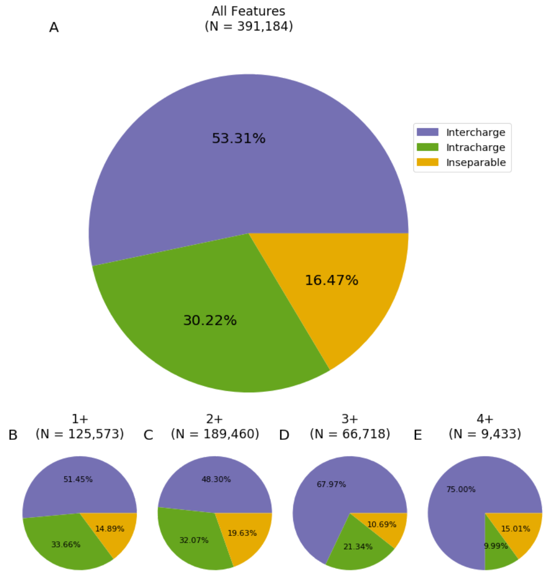


***Figure 3: Intracharge separation power is stronger than Intercharge separation power.*** *Evaluation of effects of IM on intercharge and intracharge separation for (A) average feature, and for each individual charge state (B-E). Pairwise feature overlaps are classified as separable due to intercharge influences (purple), separable due to intracharge influences (green) or inseparable by IM (yellow).*

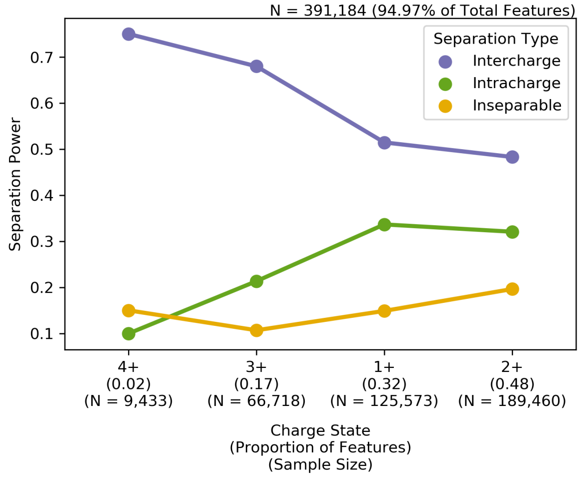


***Supplemental Figure 4 - Intercharge separation power decreases with increased proportion of peptide features in charge state.*** *Relationship between proportion of peptide features in a charge state and the proportion of overlaps classified as separable by intercharge, intracharge or inseparable. Separation power is a measure of the proportion of pairwise overlaps separable by IM in a separation type group for an average peptide in that charge state. Features with no overlaps without IM are excluded from the analysis (N = 20 739). As the proportion of features in a charge state increases, the intercharge separation power decreases and intracharge separation power increases.*

**Relationship between IM isolation window size and spectral complexity**

In targeted Data Independent Acquisition (DIA) workflows, future improvements to IM resolution and IM-based peptide libraries will increase library accuracy and thus decrease the required post-acquisition IM data extraction window. To investigate the impact of IM isolation window width on spectral complexity in a diaPASEF[2] workflow, we examined the relationship between IM width size and the average number of features isolated per window in IM widths ranging from 0.03 to 0.30 1/k0. This range captures the current IM computational extraction width of 0.06 1/K0 and the experimental isolation width of 0.25 1/k0. At IM widths below 0.06 the relationship between IM width and average number of precursors per window is almost linear ($R^2$ = 0.98) (Fig 2 inlet), suggesting that a decrease in IM width would lead to a linear decrease in spectral complexity. At IM widths around 0.25 the number of features begins to saturate. This saturation is likely due to the ability for larger IM width sizes to reliably separate features between charge states, while providing poor intracharge separation at this low resolution (Fig 2). This suggests that in a certain IM width range, the width can be changed without significant changes to the spectral complexity. These findings provide insight into the nature of IM separation in the context of a diaPASEF experiment and suggest that improved IM resolution will allow for further reduction of spectral complexity.

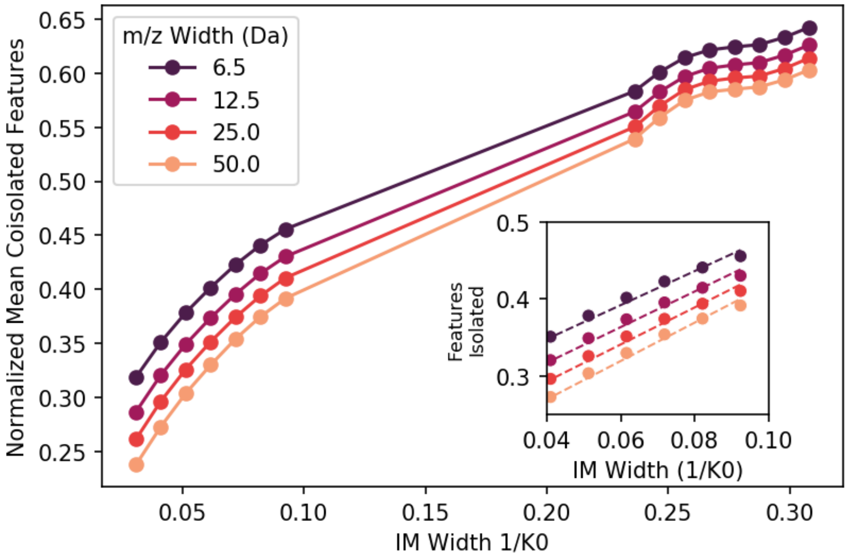


***Figure 5 Relationship between IM width coisolation rate.*** *Mean number of coisolated peptides per isolation window as a function of IM width for varying m/z widths. Means are normalized across different m/z widths by computing the proportion of coisolation events that are unresolved by IM at a given m/z and IM width. At low IM widths (0.03-0.09 1/k0) indicative of the IM extraction width in a diaPASEF workflow, the mean coisolated features per isolation window decreases linearly with decreased IM width ($R^2$ = 0.98). The number of coisolated features saturates at a larger IM width (0.23-0.30 1/k0), indicative of the experimental IM widths in a diaPASEF workflow.*

### References

1. Meier, F. *et al.* Online parallel accumulation – serial fragmentation (PASEF) with a novel trapped ion mobility mass spectrometer. *Mol. Cell. Proteomics* (2018) doi:10.1074/mcp.TIR118.000900.
2. Meier, F. *et al.* diaPASEF: parallel accumulation–serial fragmentation combined with data-independent acquisition. *Nat. Methods* **17**, 1229–1236 (2020).
